# Multiscale modelling of neuronal dynamics in hippocampus CA1

**DOI:** 10.1101/2024.04.17.589863

**Authors:** Federico Tesler, Roberta Maria Lorenzi, Adam Ponzi, Claudia Castellato, Fulvia Palesi, Daniela Gandolfi, Claudia A.M. Gandini Wheeler Kingshott, Jonathan Mapelli, Egidio D’Angelo, Michele Migliore, Alain Destexhe

**Affiliations:** Paris-Saclay University, CNRS, Paris-Saclay Institute of Neuroscience (NeuroPSI), 91198 Gif-sur-Yvette, France; Department of Brain and Behavioural Sciences, University of Pavia, Pavia, Italy; Institute of Biophysics, National Research Council, Palermo, Italy; Digital Neuroscience Centre, IRCCS Mondino Foundation, Pavia, Italy; Department of Biomedical, Metabolic and Neural Sciences, University of Modena and Reggio Emilia, Via Campi 287, 41125, Modena, Italy; NMR Research Unit, Queen Square MS Centre, Department of Neuroinflammation, UCL Queen Square Institute of Neurology, Faculty of Brain Sciences, University College London, London, United Kingdom

**Keywords:** spiking neural network, hippocampus, mean-field, traveling waves, oscillations

## Abstract

The development of biologically realistic models of brain microcircuits and regions is currently a very relevant topic in computational neuroscience. From basic research to clinical applications, there is an increasing demand for accurate models that incorporate local cellular and network specificities, able to capture a broad range of dynamics and functions associated with given brain regions. One of the main challenges of these models is the passage between different scales, going from the microscale (cellular) to the meso (microcircuit) and macroscale (region or whole-brain level), while keeping at the same time a constraint on the demand of computational resources. One novel approach to this problem is the use of mean-field models of neuronal activity to build large-scale simulations. This provides an effective solution to the passage between scales with relatively low computational demands, which is achieved by a drastic reduction in the dimensionality of the system. In this paper we introduce a multiscale modelling framework for the hippocampal CA1, a region of the brain that plays a key role in functions such as learning, memory consolidation and navigation. Our modelling framework goes from the single cell level to the macroscale and makes use of a novel mean-field model of CA1, introduced in this paper, to bridge the gap between the micro and macro scales. To develop the mean-field model we make use of a recently introduced formalism based on a bottom-up approach that is easily applicable to different neuronal models and cell types. We test and validate the model by analyzing the response of the system to the main brain rhythms observed in the hippocampus and comparing our results with the ones of the corresponding spiking network model of CA1. In addition, we show an example of the implementation of our model to study a stimulus propagation at the macro-scale, and we compare the results obtained from our model with the corresponding spiking network model of the whole CA1 area.

## 1 Introduction

The development of large-scale models and simulations of brain activity (going from thousands of neurons to full regions and whole-brain scale) has seen a great advance in the last few years, boosted by the increase of the computational power and modelling tools. Many of these models are based on relatively detailed single-cell models and data-driven connectivity structures, which allows to build simulations that can capture the specificities of local brain circuits [1–3]. Even when the advances have been remarkable, these detailed models demand high computational resources and are restricted to local circuits or brain regions, while building models at whole-brain level with single-cell resolution is still far from possible. Thus, an alternative solution that allows to move efficiently between scales (from cells to regions to whole-brain) is currently of great importance. One possibility has recently emerged which consists on using mean-field models of neuronal activity to build large-scale simulations [4–6]. Mean-field models use statistical techniques to estimate the activity of large neuronal populations (from hundreds to thousands of neurons), which allows to reduce the dimensionality of the system. Thus, the activity of local brain circuits can be described in terms of a few differential equations, which drastically reduce the need of computational resources. The low-dimensionality of these models make them very good candidates to be integrated into large-scale simulations. Recently developed computational tools, such as the The Virtual Brain, make use of mean-field field models together with connectome information to build whole-brain simulations, and which can be performed without the need of large computational resources [4]. This approach has been applied to whole-brain simulations for different species and is being used in basic research [6, 7] and for clinical applications [8, 9], which shows the relevance and utility of these methods. Although the results obtained so far are notorious, these methods are normally based on generic mean-field models (sometimes inspired on cortical microcircuits), which do not incorporate the specificities of the different brain regions. However, the different activity patterns and functions that characterize each region is intrinsically linked to the specific cell-types and local connectivity structure observed in each area. Thus, in order to extend the utility and applicability of these methods it is of fundamental importance to incorporate the cellular heterogeneity and structural specificity observed in the brain. Some attempts in this direction have been done, mostly based on phenomenological mass-models adapted to capture particular dynamics [10, 11], but which do not capture cell specificity and local connectivity structures. Only recently detailed mean-field models of a specific sub-cortical microcircuit have been proposed for the cerebellar cortex [12], thalamus [13] and basal ganglia [14]. Thus, further developments in this direction are of fundamental importance.

In this paper we introduce a multiscale model of the hippocampus which incorporates a newly developed mean-field model as the bridge between the different scales. In particular we focus on the hippocampal CA1, an area known for playing a key role in main brain functions such as learning, memory consolidation and navigation [15–17]. To develop the mean-field of the CA1 microcircuit we make use of a recently developed formalism that follows a bottom-up approach starting from the single-cell level, which allows to build a mean-field model that incorporates different cell types with specific intrinsic firing properties, and their synaptic interactions mediated by different receptor types [18–20]. In addition we develop a macroscale simulation of CA1 using the mean-field models as building blocks and incorporating extended specific connectivity structure based on a recently developed data-driven method [2] (see Fig. 1 for a diagram of the multiscale framework).

**Fig. 1.**
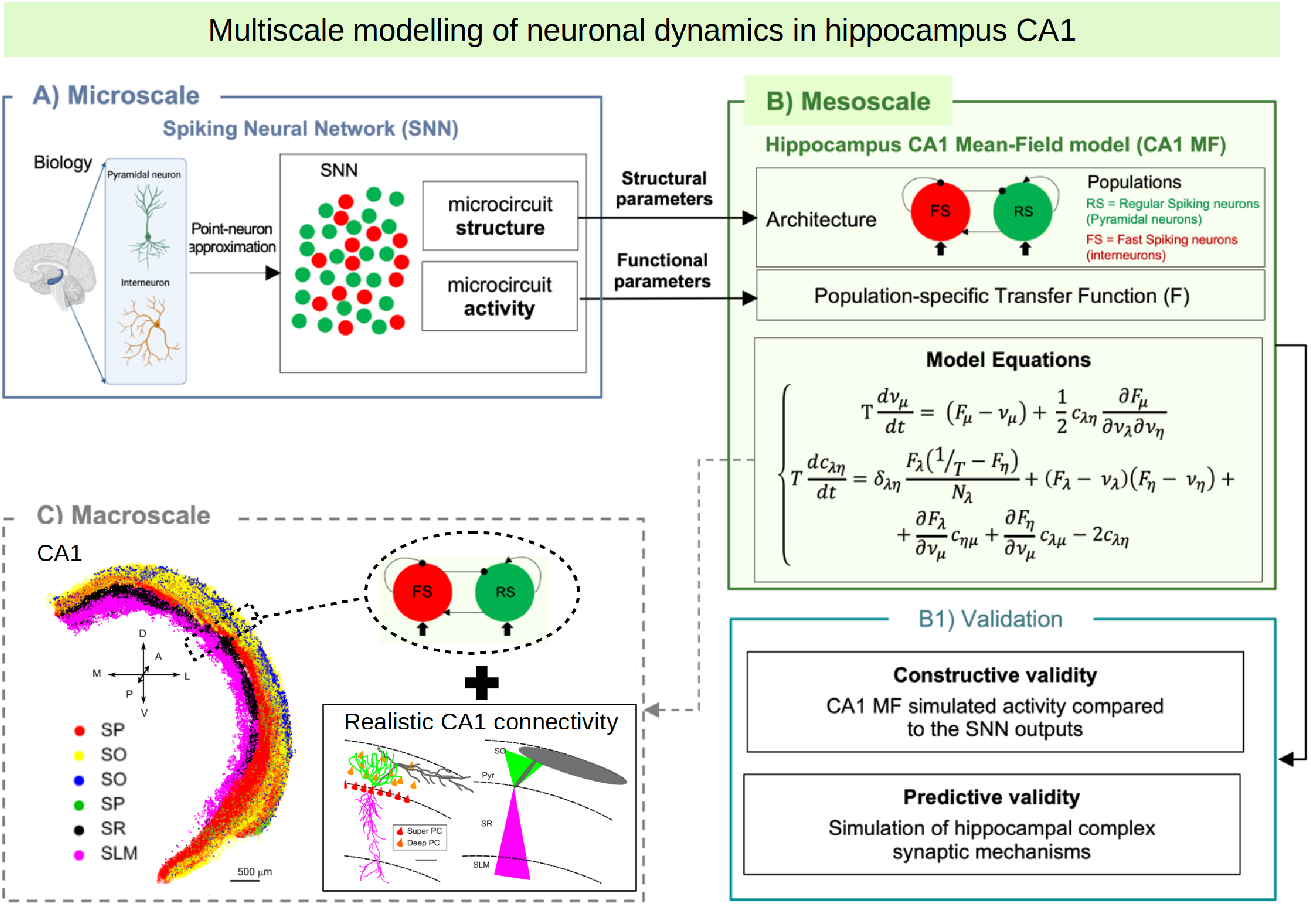
Diagram of the multiscale modeling framework. A) Starting at the single-cell level, we build spiking neural networks taking into account cellular and local connectivity properties of hippocampus CA1. B) We develop a mean-field model of the network dynamics using a recent bottom-up formalism, which incorporates the the cellular and network specificities. C) Finally we build a macroscale model using the mean-field models (representing columns or local domains) in combination with realistic extended connectivity of CA1. The image of CA1 is adapted from Ref. [2]. The color code represent different neurons in 4 layers of CA1: red: Superficial Pyramidal Cells (SP); yellow: Deep Pyramidal Cells (SO); blue: Stratum Oriens Inhibitory neurons (SO); green: Stratum Pyramidalis Inhibitory neurons (SP); black: Stratum Radiatum Inhibitory neurons (SR); magenta: Stratum Lacunosum Inhibitory neurons (SLM).

In the next sections we first present the model of the CA1 microcircuit and the mean-field formalism with more details and describe the development of the CA1 mean-field model. Then we will test and validate our model by analyzing the multiscale model response under the main oscillatory activity observed in the hippocampus and comparing the mean-field model results with to the ones of an equivalent spiking network model. Then we will show how the mean-field model can be used to build a macroscale simulation taking into account the realistic extended connectivity of CA1. The modelling framework presented here allows us to go from single-cell models to biologically realistic macroscale simulations while keeping a limited use of computational resources. In addition, the model is suitable to be incorporated into whole-brain simulation platforms (such as the TVB [4]), which highlights the importance and usability of this approach.

## 2 Methods

### 2.1 Single-cell model

Our multiscale modelling starts at the single-cell level. To perform single-cell simulations we adopt the Extended-Generalized Integrate-and-Fire neuronal model (EGLIF) [12, 21]. The equations for the EGLIF model describe the time evolution of membrane potential (*V*_*m*_), slow adaptation current (*I*_*adap*_) and fast depolarization current (*I*_*dep*_):

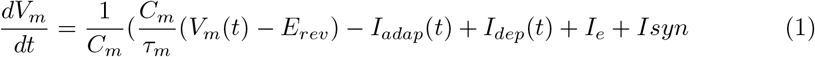

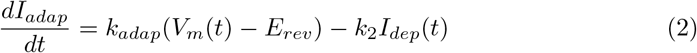

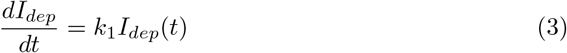

where *I*_*syn*_ is the synaptic current modeling the synaptic stimulus, *C*_*m*_ is the membrane capacitance, *τ*_*m*_ is membrane time constant, *E*_*rev*_ is the reversal potential, *I*_*e*_ is the endogenous current, *k*_*adap*_ and *k*_2_ are adaptation constants and *k*_1_ is the decay rate of *I*_*dep*_. When a spike occurs at time *t*_*spk*_, the update rules of the state variables is given by:

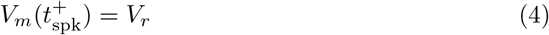

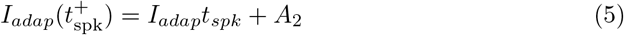

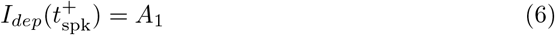

where 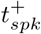 is the time instant immediately following *t*_*spk*_, *V*_*r*_ is the reset potential, and *A*_1_ and *A*_2_ are the model currents update constants. For our simulations we will consider only two types of cells (pyramidal cells and fast spiking interneurons (FS)), although the model could be extended to incorporate more cell types. We note that a data-driven adaptive GLIF model (AGLIF) has been recently developed [22], specifically conceived to capture the detailed dynamics observed experimentally in CA1 neurons and interneurons. In this work, we used a simplified EGLIF implementation, which is more easily adaptable to the multiscale formalism introduced in this paper while still provides an effective way of simulating the neuronal and population dynamics as will be shown in the next sections. The model parameters used for each cell type are given in 1. It should be stressed that the mean-field formalism used for the analysis in the following sections has shown to be robust for large variations in neuronal parameters [20, 23], for which the specific values used here serve as a general reference for building our system.

**Table 1.** Neuronal parameters for the EGLIF model. We consider two cell types, pyramidal neurons (Pyr) and fast spiking inteneurons (FS).

|  | Pyramidal cells (Pyr) | Interneurons (FS) |
| --- | --- | --- |
| $C_m$ (pF) | 2877.83 | 2939.66 |
| $\tau_m$ (ms) | 10955.36 | 2169.40 |
| $E_{rev}$ (mV) | -70.07 | -74.01 |
| $k_{adap}$ (MH<sup>-1</sup>) | 0.0084 | 0.0616 |
| $k_1$ (kHz) | 0.0007 | 0.0021 |
| $k_2$ (kHz) | 0.0042 | 0.0098 |
| $A_1$ (pA) | 26.0 | 92.0 |
| $A_2$ (pA) | 170.0 | 5.0 |
| $I_e$ | 0 | 0 |

### 2.2 CA1 microcircuit and Mean-Field formalism

The second scale of our modelling framework is at the microcircuit level. For simplicity we will assume that the circuit is made of two cell-types, pyramidal excitatory cells (Pyr) and fast spiking inhibitory interneurons (FS), where each cell will be modeled with an E-GLIF model presented in the previous section. For the initial construction of the model we will consider a network of 5000 Pyr-cells and 500 FS-cells. Neurons in the circuit are interconnected with probability *p*_*Pyr*−*Pyr*_ = 0.01, *p*_*FS*−*Pyr*_ = 0.3, *p*_*Pyr*−*FS*_ = 0.2, *p*_*FS*−*FS*_ = 0.3 [24, 25]. The local microcircuit receives external excitatory input from the CA3 area, which will be modeled as an external poissonian input representing 5000 excitatory neurons. The external input targets both Pyr and FS cells with probability of *p*_*ext*−*Pyr*_ = 0.15 and *p*_*ext*−*FS*_ = 0.3 respectively [25].

Next, we introduce the mean-field model of the CA1 miorcircuit dynamics. To develop this mean-field model we will adopt a recent formalism adapted for EGLIF neurons. The formalism is based on a bottom-up approach, starting at single-cell level, which allows the construction of mean-field models with cellular-type specificity. The mean-field equations for the E-GLIF network are given to a first-order by [12]:

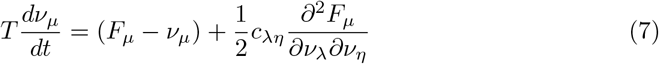

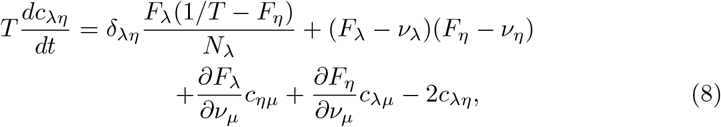

where *ν*_*j*_ is the mean neuronal firing rate of the population *j, F* is the neuron transfer function (i.e. output firing rate of a neuron when receiving the corresponding excitatory and inhibitory inputs with mean rates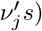, and *T* is a characteristic time for neuronal response (we adopt *T* = 5 ms).

Following Zerlaut et al [19] we write the transfer function for each neuronal type as:

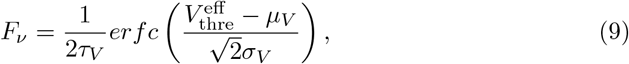

where *erfc* is the error function, 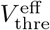 is an effective neuronal threshold, *µ*_*V*_, *σ*_*V*_ and *τ*_*V*_ are the mean, standard deviation and correlation decay time of the neuronal membrane potential. The effective threshold can be written as a second order polynomial expansion:

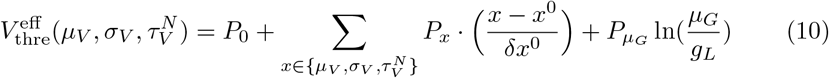

where *x*^0^, *δx*^0^ are constants, the coefficients *P*_*x*_ are to be determined by a fit over the numerical transfer function obtained from single-cell spiking simulations for each specific cell-type, and where *µ*_*G*_ is given by:

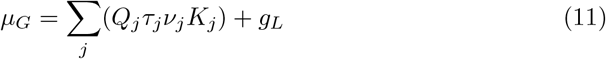

with *K*_*j*_ the mean synaptic convergence of type *j*.

We can write the mean membrane potential and standard deviation as [12]:

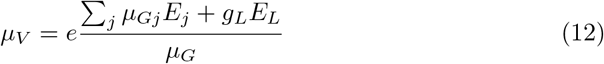

Finally, the standard deviation and correlation decay time of the neuronal membrane potential can be written as:

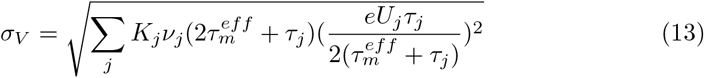

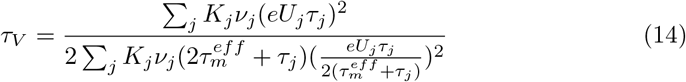

With 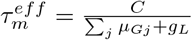 and 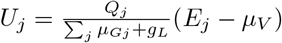.

The derivation of the mean field equations can be found in Refs. [20] and [12].

## 3 Results

We start by deriving the mean-field model of CA1, then we compare this model with spiking network simulations, for different situations, and we terminate by showing mesoscale phenomena such as traveling waves in large-scale systems.

### 3.1 Mean-Field model of CA1 microcircuit

To build the mean-field model for the hippocampus we first calculate the corresponding transfer function for each cell type. This is done by fitting the numerical transfer function obtained from single-cell simulations to Eq.10. In Figure. 2 we show the numerical transfer function together with the fit for each cell-type.

**Fig. 2.**
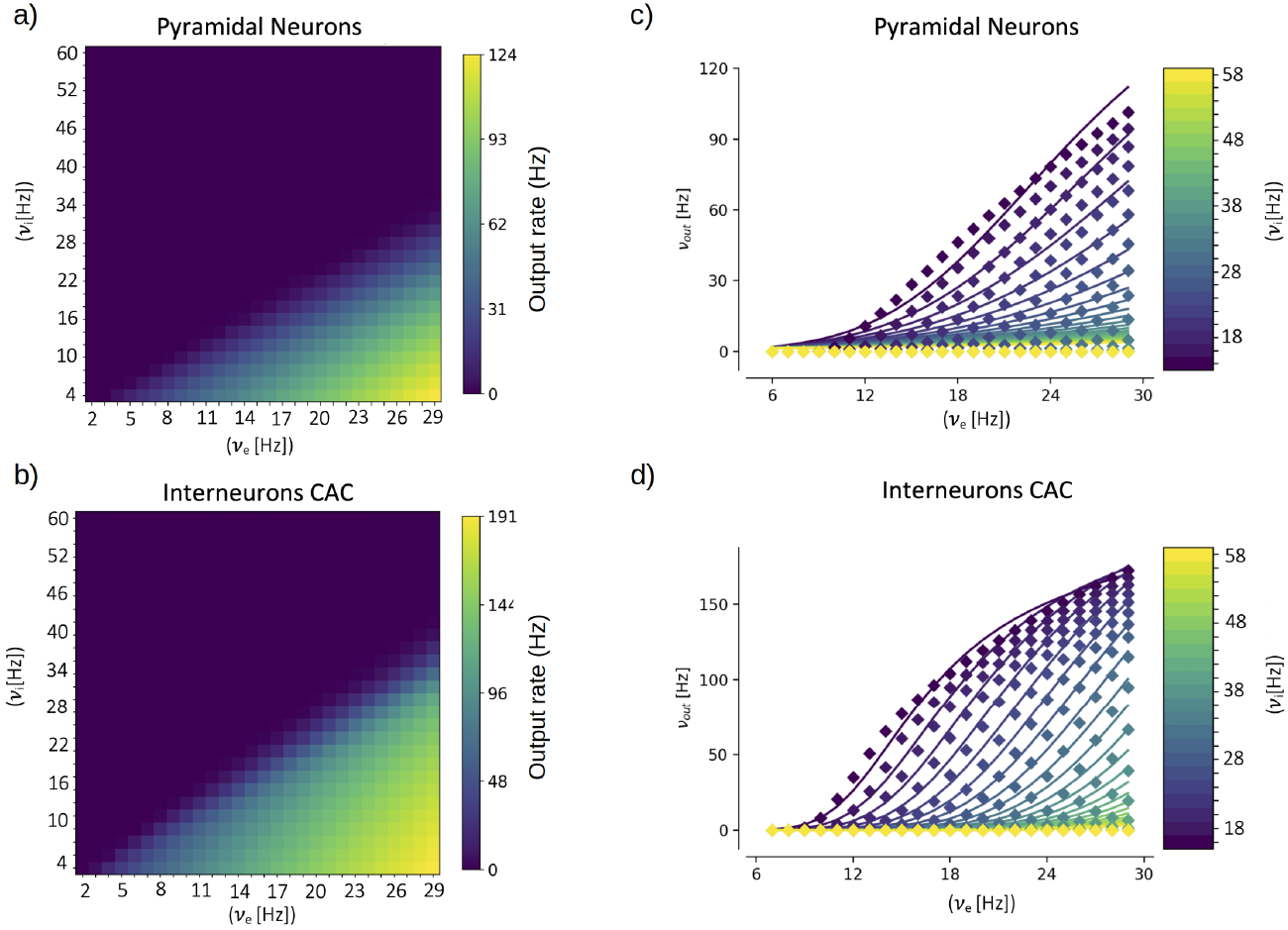
Numerical transfer function (a,b) and the corresponding semi-analytical approximation fitted from Eq.9 (c, d) for each cell-type. Solid lines in panels c,d correspond to the firing rates obtained from Eq.9 while filled-squares correspond to the numerical results.

### 3.2 Activity patterns and time varying inputs

To validate the multiscale model of the hippocampus we test the response of the model to some of the main activity patterns observed in CA1. It is well established that three main patterns of activity are present in the hippocampus, and can be observed during specific brain states: theta oscillations (4-10Hz) are normally associated with exploratory behaviour, sharp-wave/ripple complexes (140-200 Hz) are associated with immobility, and gamma oscillations (40-140 Hz) are normally present in combination and modulated by the other two rhythms. In Fig. 3.a and b we show results of the simulation for stimulations on the theta and gamma ranges. We show the results obtained with mean-field superimposed to the results from the spiking neural network (SNN). As we can see the mean-field can correctly reproduce the response of the system for the different input patterns.

**Fig. 3.**
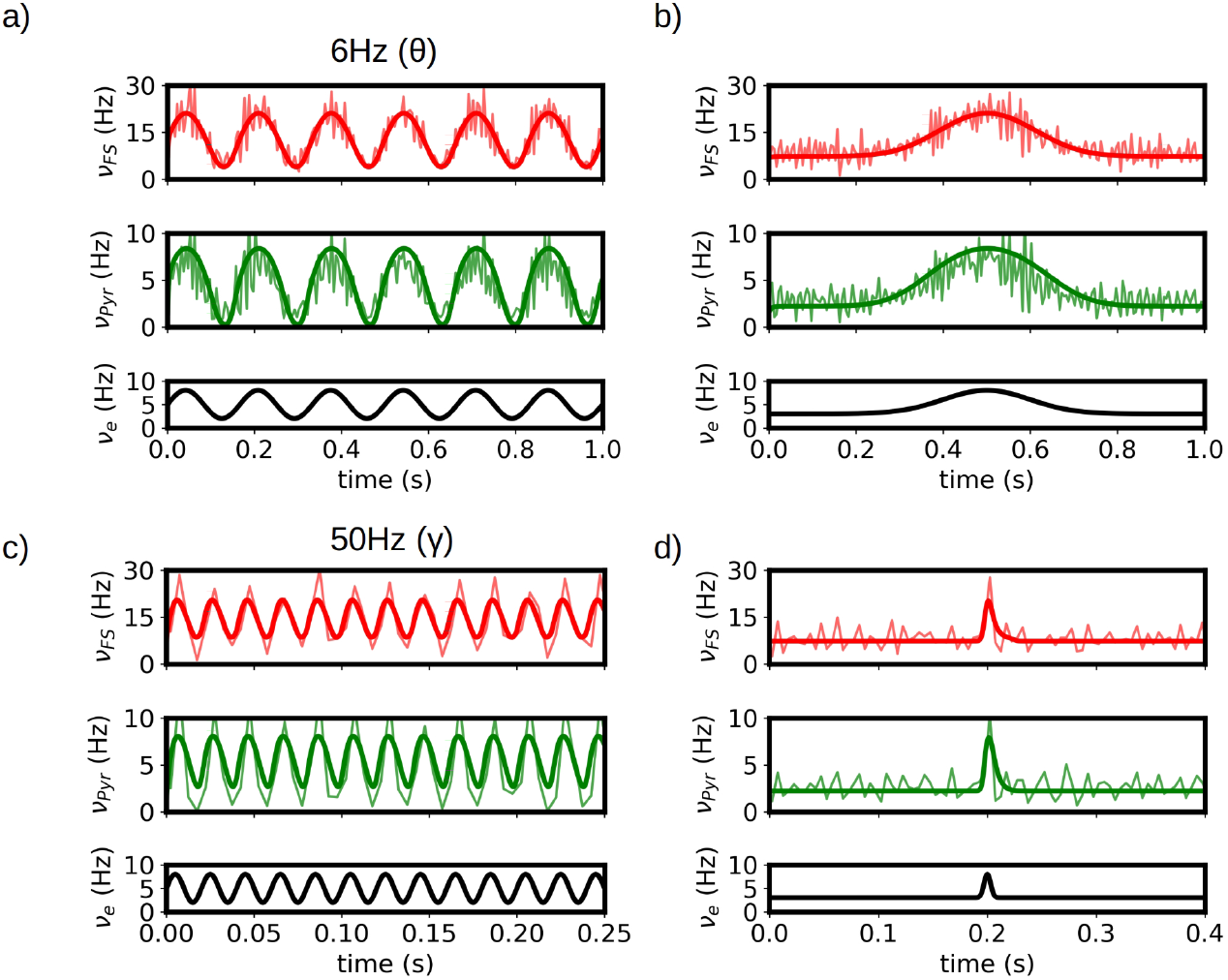
Response of the systems to *θ* (a) and *γ* (b) rhythms. Results from the mean-field (bold solid lines) are superimposed to the firing rates obtained from the spiking-network (SNN) simulations of the hippocampus (light solid lines). c,d) Response of the system to slow and fast Gaussian inputs. We see that the mean-field can capture the response of the SNN in large frequency-range (from 6Hz in *θ* waves to ∼140Hz for the fast Gaussian input), relevant to simulate the different activity patterns observed in the hippocampus. For high frequency the accuracy of mean-field is reduced as the typical time of variation in activity gets closer to the characteristic times of the mean-field.

In addition, in Fig. 3.c and e we show the response of the system to low and fast Gaussian-shaped inputs. The fast input can be seen as similar to the activity of sharp-waves in CA1, while the slow input can be seen as a typical response curve of place cells in CA1 for space-field selectivity. The mean-field is capable of capturing the response of the system for both cases. For fast or high-frequency inputs the accuracy of mean-field is slightly reduced as the typical time of variation in activity gets closer to the characteristic times of the mean-field.

### 3.3 Synaptic potentiation and depression

The occurrence of long term synaptic depression (LTD) and potentiation (LTP) in the hippocampus was among the first experimental studies presented on long term synaptic plasticity and is believed to be related with the role of hippocampus in learning and memory formation, one of the main known functions of this region [26, 27]. The capacity of reproducing changes in the synaptic strength is a key feature to be captured by a model of this region. To perform this study we analyze the response of our mean-field model under variations in the synaptic convergence. In particular we consider variations in the synaptic convergence of the simulated CA3 afferent input to the local Pyramidal cells in CA1 (see diagram in Fig. 4.a). We introduce the parameter *W*_*e*_ which quantifies the changes in the weight of the synaptic convergence, being *W*_*e*_ = 100% the baseline level (as considered in the previous sections), and we analyze the response of the system for a variation of 50% in the strength of the synaptic convergence for a constant input and a time varying input. In Fig. 4.a we show the evolution of the response of pyramidal cells as a function of *W*_*e*_ and its comparison with the results from the spiking neural network. We can see that, although there is a small overestimation of the activity for certain values of *W*_*e*_, the mean-field model can correctly capture the evolution of the response obtained in the spiking network.

**Fig. 4.**
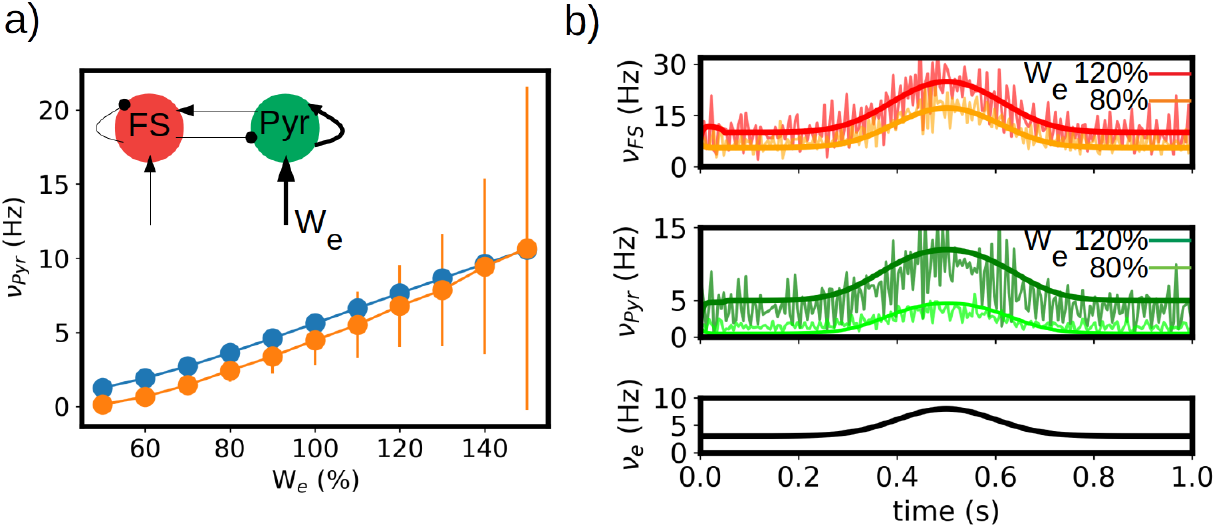
Synaptic potentiation and depression in the mean-field model. a) Evolution of the response of pyramidal cells as a function of the strength of synaptic convergence (*W*_*e*_). We show the results obtained for the mean-field model (blue circles) and the spiking neural network (orange circles) for a constant external input of *ν*_*e*_ = 5Hz. A level of *W*_*e*_ = 100% correspond to the baseline level. Inset: diagram of the network and indication of the change in convergence. b) Time varying inputs for two levels of *W*_*e*_. We show the firing rates of the FS and Pyr cells obtained from the mean-field and the spiking network together with the applied input (*ν*_*e*_).

In Fig. 4.b we show the response of the mean-field and spiking network under a timevarying input of Gaussian shape for two different levels of *W*_*e*_ (*W*_*e*_ = 80% and 120%). As we can see the mean-field can correctly reproduce the response of the network for the different values of *W*_*e*_.

### 3.4 Detailed connectivity structure and macro-scale simulations of the CA1 network

In this section we show an example of the passage from the mesoscale to the macroscale with the use of the mean-field model. As discussed before, one of the main goals of our approach is to build a model of a specific area with realistic connectivities based on available physiological, morphological and anatomical data. In this section we will present the results of simulations of a network representing a slice of hippocampal CA1 area. To this end we will adopt a recently developed method to incorporate realistic morpho-anatomical connectivities based on the geometrical probability volumes associated with pre- and postsynaptic neurites [2]. The method has been benchmarked for the mouse hippocampus CA1 area, and the results show that this approach is able to generate full-scale brain networks that are in good agreement with experimental findings. Following Gandolfi et al [2], we will focus on a particular case where only excitatory connections are taken into account, a case which has been previously compared to experimental results [2]. In Fig.5.a-b) we show a diagram of the geometric probability volume associated with pyramidal cells and the distribution of Pyr cells in CA1, adapted from Ref.[2]. We will assume that the Pyr cells are homogeneously distributed over the Pyr and SO layers. The geometric probability volumes associated with the basal, apical dendrites and axon are indicated in green, pink and grey respectively. Axonal volumes can be represented by a combination of two elliptical volumes, while dendritic volumes can be represented by conical volumes. The most relevant region for Pyr-to-Pyr connectivity lies within the Pyr-SO region, we will therefore concentrate our attention on this area to build our network. We will consider a slice covering a surface of 1.5×1.5 mm^2^ along the Pyr-SO layer. We will divide this area in compartments of 100*µm*x100*µm* containing about 200 neurons each and we will describe each of this compartment with a single mean-model as described in the previous sections. To build the connectivity between compartments we will make use of the geometric probability volumes. In Fig.5.c) we show a diagram of the compartmentalization and the corresponding single-cell probability volumes. The connectivity between compartments (given by the parameter *K* in Eq.11) will be defined as proportional to the normalized probability of connections given by the probability volumes. Here we assume that the dendritic volumes extend through the entire transverse length of the Pyr-SO layer, for which we assume that the main constraint for the connectivity is given by the axonal volume (see Fig.5).

**Fig. 5.**
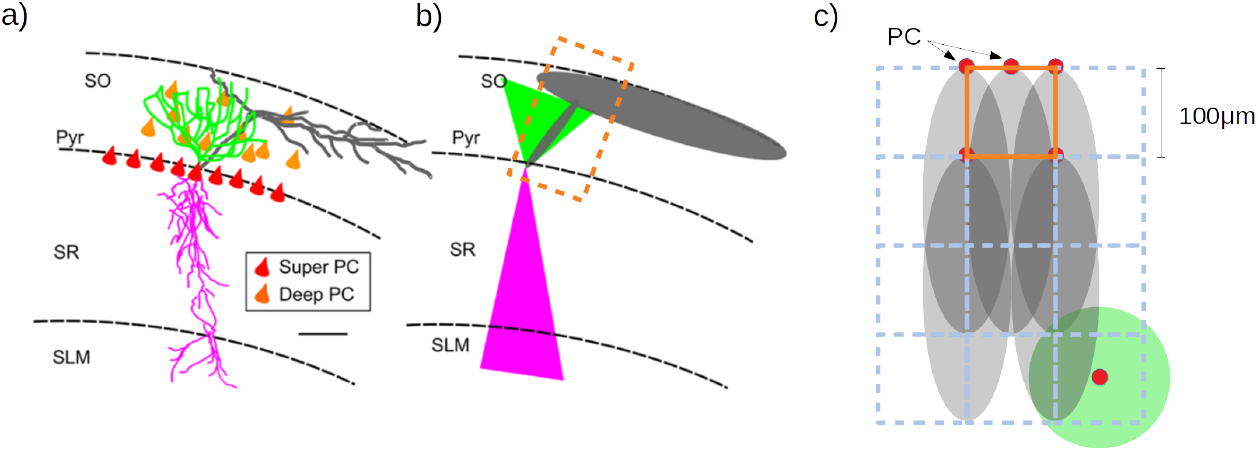
a) Realistic morphology of a superficial pyramidal cell (PC) with basal dendrites in green, apical in pink, and axon in gray, oriented within a region of a transversal CA1 hippocampal slice. Red triangles correspond to PC soma location within the stratum pyramidalis whereas orange triangles represent the scattered distribution of deep PCs within the SO. b) Probability clouds of connectivity represented as two triangles (2D of a cone) and an ellipse (2D of an ellipsoid). Color code respects the realistic morphology. The dashed rectangle in dark-orange corresponds to the area covered by a single mean-field compartment described in panel (c). c) Diagram of the compartmentalization for the mean-field description of the hippocampal network and the corresponding single-cell probability volumes (top view). Color code respects panels (a) and (b). Axonal probability clouds are shown for 5 pyramidal cells (with somas indicated in red-circles) located at the border of a compartment (indicated in dark-orange). Neighboring compartments are shown in dashed blue lines. Probability cloud for basal dendrites of single PC cell is shown at the bottom right with the soma located at the center of the compartment (red circle). Panels (a), (b) are adapted from Ref.[2]

It has been shown experimentally that in the absence of synaptic inhibition CA1 activity shows strong directionality from the CA3 side to the subiculum side. This has been also reproduced by spiking network simulations of CA1 following the same geometric connectivity volume approach. To validate our network we show in Fig.6 the results from the mean-field network slice together with the results from the corresponding spiking network simulation. In this simulations a short stimulus is applied to a single compartment in the case of the mean-field and to ∼ 200 neurons close to the CA3 region. As we can see the connectivity profile induces a strongly directed propagation from the CA3 to the Subiculum direction. In addition, the propagation evolves with an increase in neuronal recruitment which in turns leads to the appearance of a lateral propagation as the activity gets closer to the CA1-Subiculum edge. These two features can be well captured by the mean-field network.

**Fig. 6.**
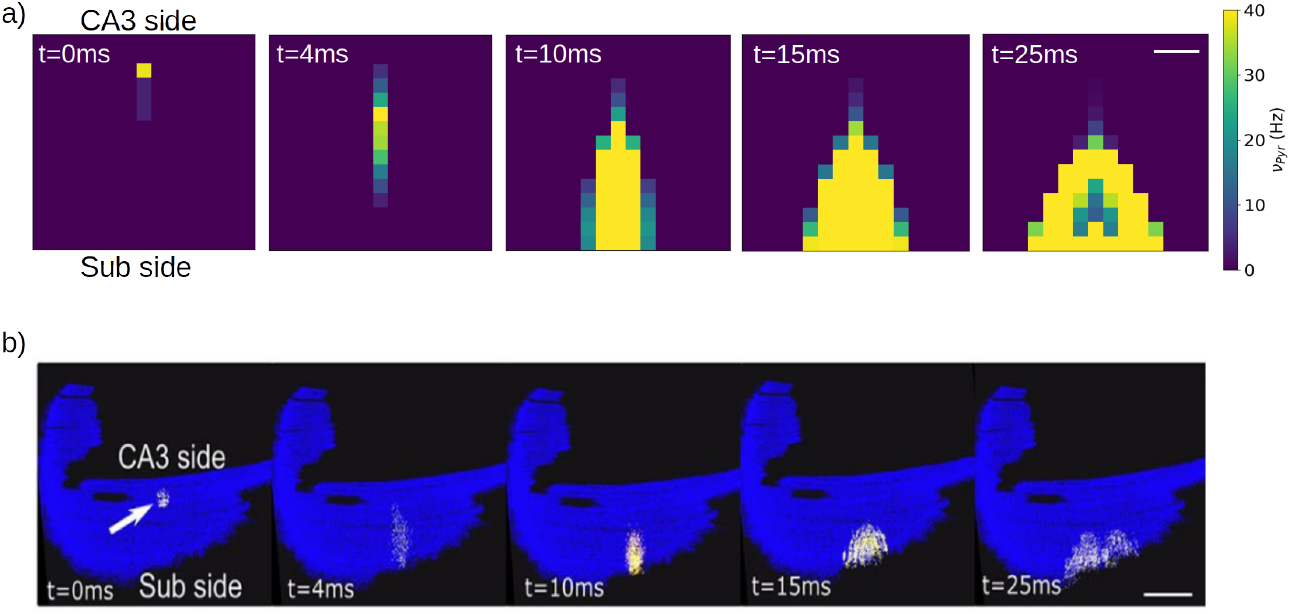
Simulation of a local stimulation in a CA1 network. Activity is evoked near the CA3 side in area of 1e4*µ*^2^ containing approximately 200 pyramidal neurons, represented by a single meanfield model. The stimulation induces a rapid propagation of the activity in the transversal direction (antero-posterior) of the network (4,10ms) with a gradual increase in neuronal recruitment and a subsequent propagation in the longitudinal direction (medio-lateral). The network correspond to a slice of 1.5×1.5 mm. Firing rates are indicated on the colorbar. Scale bar 300*µm*. b) Stimulation protocol equivalent to (a) performed in a full CA1 spiking network, adapted from Ref.[2]. Scale bar 1 mm. Activity is color coded from blue (rest) to white (spike), to visualize action potentials, with a fixed 2 ms transition time.

## 4 Discussion

In this paper we have introduced a multiscale modelling framework of the CA1 microcircuit, which goes from the single-cell to the macroscale level. This framework incorporates a newly developed mean-field model that allow us to perform an efficient passage between the different scales. The mean-field model was built using a recently introduced formalism that follows a bottom-up approach, starting at the single-cell, which made possible to incorporate cellular and synaptic specifities of CA1 within the mean-field formulation. The single-cell parameters were based on previous detailed modeling of CA1 pyramidal neurons and fast-spiking interneurons [22], and synaptic connectivity information was based on experimental data [24, 25]. We have tested the model by analyzing the response of the model under different oscillatory rhythms found in CA1 and we have validated the results by comparison with the corresponding spiking network model. We have shown that the mean-field can correctly reproduce the results of the spiking-network for activity patterns related with some of the main patterns observed in CA1 (theta oscillations, sharp-waves and gamma oscillations).

Finally we have shown an example of the implementation of a macroscale simulation within our framework. In particular we built a simulation of a slice of CA1 with specific connectivity structure, based on a recently developed data-driven method [2]. Furthermore, we compared the results of our simulations with an equivalent simulations of a spiking-network model of CA1, showing that our model can capture some of the main features of the spiking simulations, which further validates our model.

The model presented in this work is a step forward to the development of regionspecific multiscale models. In addition, the framework developed here is suitable to be included in whole-brain simulation platforms [4], which extends the importance and utility of our study. Furthermore, methods to estimate brain signals (LFP, EEG, MEG, fMRI) from the type of mean-field used here have already been developed [28, 29], which will also allow the comparison with experimental results on whole-brain activity. In combination, these developments provide an efficient solution to the complicated task of modeling the brain at different scales and open new perspectives for future studies.

